# Unveiling mechanical interactions between cell division and extracellular matrix in human colonic epithelium organoids: A 4D study using DVC

**DOI:** 10.1101/2024.10.07.617033

**Authors:** L. Magne, T. Pottier, D. Michel, J. Laussu, D. Bonnet, L. Alric, G. Recher, S. Segonds, F. Bugarin, A. Ferrand

## Abstract

Cell division is a major event in tissue homeostasis, enabling renewal and regeneration. Stem cells, in particular, play an important role in this homeostasis, thanks to their ability to perform symmetric or asymmetric cell divisions. To study cell division, the human colon epithelium represents a model of choice due to its rapid renewal and therefore high proliferative potential. Currently, studying the live mechanical interactions between the epithelium and its matrix *in vivo* is challenging due to the lack of suitable methods. 3D human colon organoids seeded in Matrigel® are good models for this purpose as, from isolated stem cells, they recapitulate the tissue architecture organization and properties. This culture set-up also allows to study the matrix displacements around the organoid.

Here, we studied the impact of cell division within the colonic epithelium on the extracellular matrix. We performed and validated an original experimental and analytical process with a 3D time-lapse confocal microscopy to follow cell mitosis and matrix movements on which we performed Digital Volume Correlation. We showed that these two different types of cell division impact the matrix differently with the asymmetric divisions causing a mainly uniaxial displacement, whereas symmetric ones involved a multiaxial and more important one.

## Introduction

Cell division is a crucial process in tissues as it is involved during development, homeostasis as well as in regenerative process ^1^. Once adult, the epithelia, which represent the boundaries between our inner body and the outside world, display high number of cell division events to assure their renewal and maintain their barrier function. Among them, the colonic epithelium acts as a selective barrier between the content of the intestinal lumen (food bolus, microbiota, mucus…) and the lower layers of the colonic mucosa. This epithelium consists in a monolayer of columnar cells lining two distinct compartments, the crypts that form invaginations and the plateaus corresponding to flat areas. This epithelium is among the most proliferative and fastest-renewing one of our body assuring the renewal of the entire intestinal lining within a week ^2^. It finds its origin at the bottom of the crypts which shelter the intestinal stem cells (ISC) assuring the renewal of the epithelium. Extensive proliferation occurs within the crypt compartment due to the stem cells and the transient amplifying cells, corresponding to progenitors, which will differentiate when reaching the plateau to generate the lineages of differentiated cells assuring the functions of the tissue (water and ions absorption, protection against pathogens…).

The intestinal epithelium renewal is precisely controlled and depends on the spatial organization of signals from the crypt environment, namely the intestinal niche. In physiological conditions, the intestinal niche controls the homeostasis of the epithelium and the integrity of the crypt ^3^. This niche includes the extracellular matrix (ECM) which participates to the cell fate ^4^. The dynamics of the ECM is a key element in the evolution of tissue architecture, participating to the establishment and maintenance of stem cell niche, cell differentiation but also branching or wound repair ^5^. Its biochemical properties, such as its components or growth factors and cytokines gradients, have been studied since a long date, however the last decade has seen considerable interest in understanding the effects of its mechanical interaction with the epithelium ^4^. Indeed, cells perceive physical stimuli from the matrix notably via interactions between their cell adhesion molecules linked to their cytoskeleton such as integrins ^6,7^ or ion channels such as Piezo ^8^. The epithelium can also sense the mechanical properties of the ECM (topography, stiffness, elasticity) and translate them into intracellular messages to control a range of physiological processes regulating cellular phenotypes and behaviours ^9^. Reciprocally, the epithelium itself applies mechanical loads to the matrix, notably radial and/or orthoradial tensile forces, as a result of biological processes involved in homeostasis ^10–12^. Among these processes, the contribution of stem cell and progenitor divisions are major given the high proliferation rate of these cells in the colon. Regarding their mode of division, both cell types can proliferate by symmetric division, assuring their expansion during tissue regeneration process ^13^. However, only stem cells have the capacity to self-renew indefinitely by asymmetric division, leading to their own renewal while giving birth to a progenitor cell ^14, 15,16^. Progenitors, on the other hand, have a very limited capacity for self-renewal, and after division cannot maintain their undifferentiated state indefinitely. They can divide several times by symmetrical division, but eventually differentiate. In the case of symmetric division, the mitotic spindle is mainly parallel (0-30°) to the matrix support, while in asymmetric division, the mitotic spindle will be mainly perpendicular (60-90°) to the support ^17^. In consequence, both types of division do not exert the same type of cell-matrix interrelationship, and from a biophysical perspective may strain the ECM differently ^18^.

To date, while the consequence of the mitotic spindle orientation during the cell division on cell fate determination, tissue organization, or development is increasingly better understood ^19,20^, how it affects the extracellular matrix remains largely unexplored. A main explanation is the lack of biological model allowing to combine both live cell division and ECM displacement observation in three dimensions. Considering the proliferative capacity of the colonic epithelium, the human colonic three dimensions (3D) organoid model embedded into Matrigel® represents an appropriate model to address this question. Indeed, human colon organoids are established from crypts isolated from patient’s colon biopsie that include intestinal epithelium stem cells. Cultured in 3D in Matrigel®, playing the role of their supporting matrix, these organoids display the different tissue cell populations organized in a monolayer of epithelial cells ^21^. As a consequence, they display both asymmetric and symmetric divisions as illustrated in Figure 1. Moreover, their *in vitro* culture permits their live observation by microscopy, allowing the follow-up of the tissue culture evolution.

**Figure 1.**
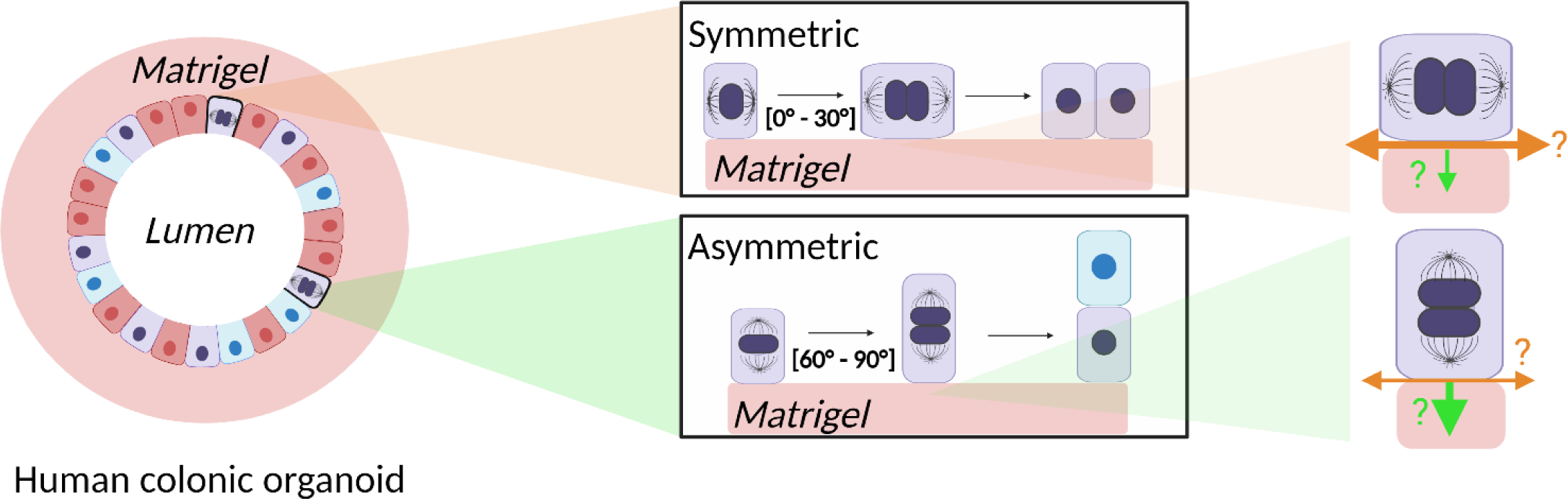
Stem cell divisions. Human colonic organoids are used as biological model to study the mechanical interactions between the epithelium and its extracellular matrix including during mitosis. Stem cell (purple on the scheme) division can be symmetric (i.e. parallel to the ECM) or asymmetric (i.e. perpendicular to the ECM). Differentiated cells are represented in blue (secretory lineage) and red (absorbent lineage). Orange arrows represent orthoradial traction and green arrows represent radial one. The width of arrows corresponds to the hypothetical importance of the traction.

In this study, we hypothesized that asymmetrical and symmetrical divisions involving different orientations of the mitotic spindle relative to the matrix will have a different impact on the 3D deformations of the surrounding ECM. Thus, the present study aims to investigate over time the impact of either asymmetric or symmetric divisions on 3D displacements of the ECM. The human colon 3D organoid model embedded in Matrigel® was used and a framework based on confocal imaging, 3D image reconstruction, cell segmentation and Digital Volume Correlation (DVC) on the matrix was developed to evaluate this hypothesis.

## Results

As aforementioned, we chose the human colon 3D organoid model to study the impact of the mitotic spindle orientation on the extracellular matrix displacements. To this end, it was essential to be able to monitor the cell divisions and the ECM displacements over time. Regarding cell division, DNA and tubulin, which forms the mitotic spindle, were visualized using Hoechst (blue channel) and a BioTracker 488 Green Microtubule Cytoskeleton Dye (green channel) respectively (figure 2a). Matrix displacement was investigated by adding fluorescent beads (red channel) into the Matrigel® (figure 2a). The organoid culture evolution was followed by image acquisition every twenty minutes over 3 hours using the Opera Phenix™ confocal microscope. We then performed 3D reconstruction of the acquired images for each of the three channels (figure 2a) and analysed firstly the mitotic spindle orientation and secondly the displacements of the nuclei as well as the matrix displacements.

**Figure 2.**
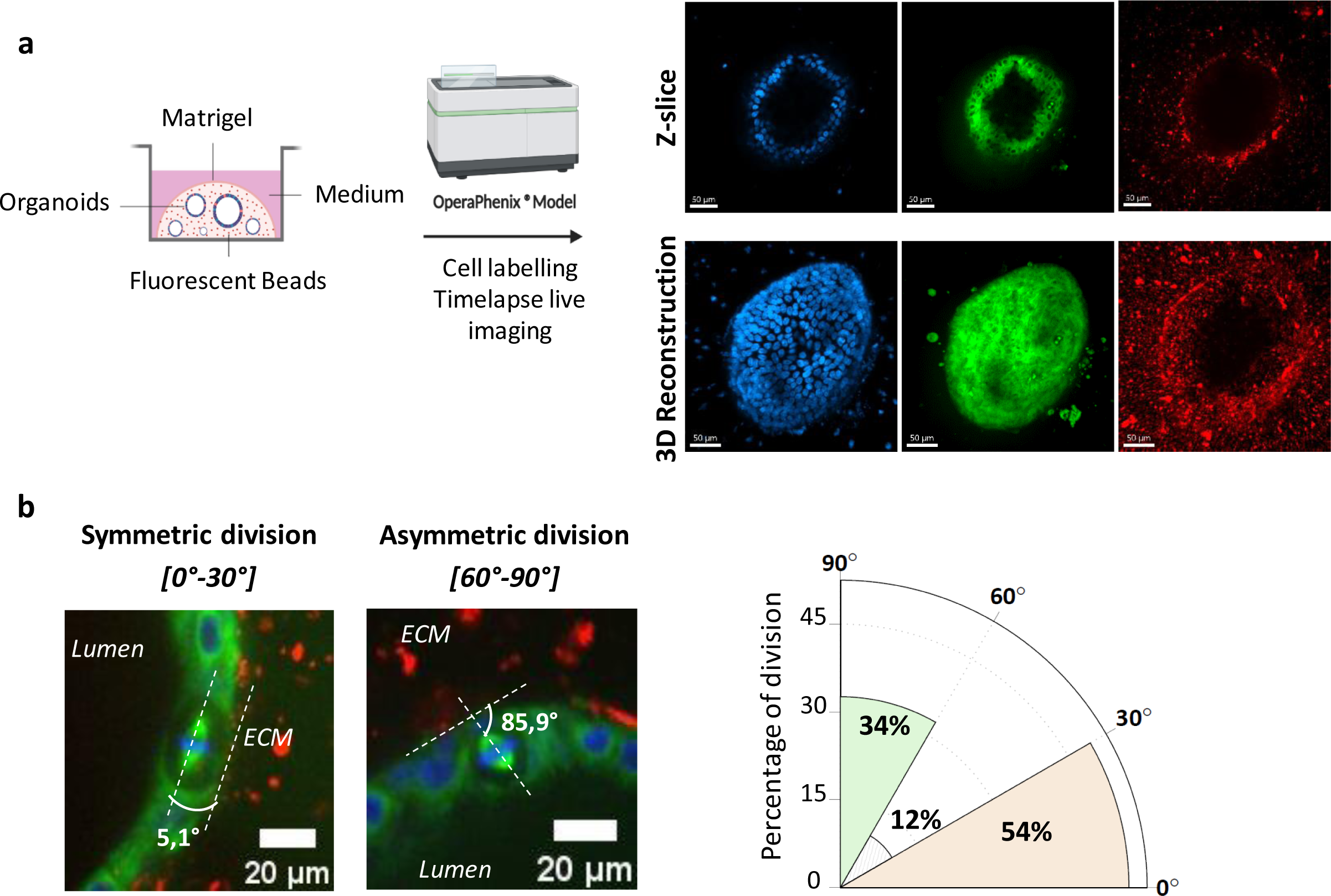
Validation of the biological model (**a**) Imaging process. Organoids are imaged with the Opera Phenix® microscope (Perkin Elmer). Nuclei are labelled with Hoechst, laser 405 nm (blue lut), tubulin is labelled with Biotracker Tubulin, laser 488 nm (green lut) and beads are imaged with laser 561 nm (red lut). (**b**) On the left, representative images of symmetric and asymmetric divisions. On the right, polar representation of the mitotic spindle orientations distribution (n=83 cells divisions from 30 organoids from 5 patients).

### Quantification of asymmetric and symmetric divisions in the human colon 3D organoid model

We first confirmed that both symmetric and asymmetric divisions occur over our epithelial organoid culture (figure 2b).

We measured the orientation of mitotic spindle during the cell divisions occurring in the human colon organoids over the three hours of observation. Mitotic spindle orientation was determined as the angle formed between the mitotic spindle axis and the organoid-Matrigel® boundary (figure 2b). In the case of asymmetrical divisions, the mitotic spindle is oriented perpendicularly, or radially, to the Matrigel. Conversely, in the case of symmetrical divisions, the spindle orientation is parallel, or orthoradial, to the support. Based on the literature ^17^, we chose to classify the angles measured into 3 intervals: [0°-30°], [30°-60°] and [60°-90°]. From a total of 83 observed cell divisions visualized in 30 different organoids of 4 different patients (a minimum of 4 organoids/patient and a maximum of 9 organoids/patients were analysed), we counted 45 mitoses with a mitotic spindle angle between 0° and 30°, i.e. 54% of divisions clearly classified as asymmetric. Mitotic spindle angle between 60° and 90°, defining symmetric divisions, were 28, i.e. 34% of divisions. It is also noticeable that a small fraction of mitoses (12%) cannot be assigned to any of the asymmetrical or symmetrical categories, with a mitotic spindle angle between 30° and 60°.

These data also demonstrate the relevance of our biological model for the study of mitotic spindle orientation.

### Impact of asymmetric and symmetric divisions on neighbouring cells

We investigated whether asymmetric and symmetric divisions might affect the neighbouring cells by focusing on the positioning of neighbouring nuclei in relation to the dividing cell (figure 3).

**Figure 3.**
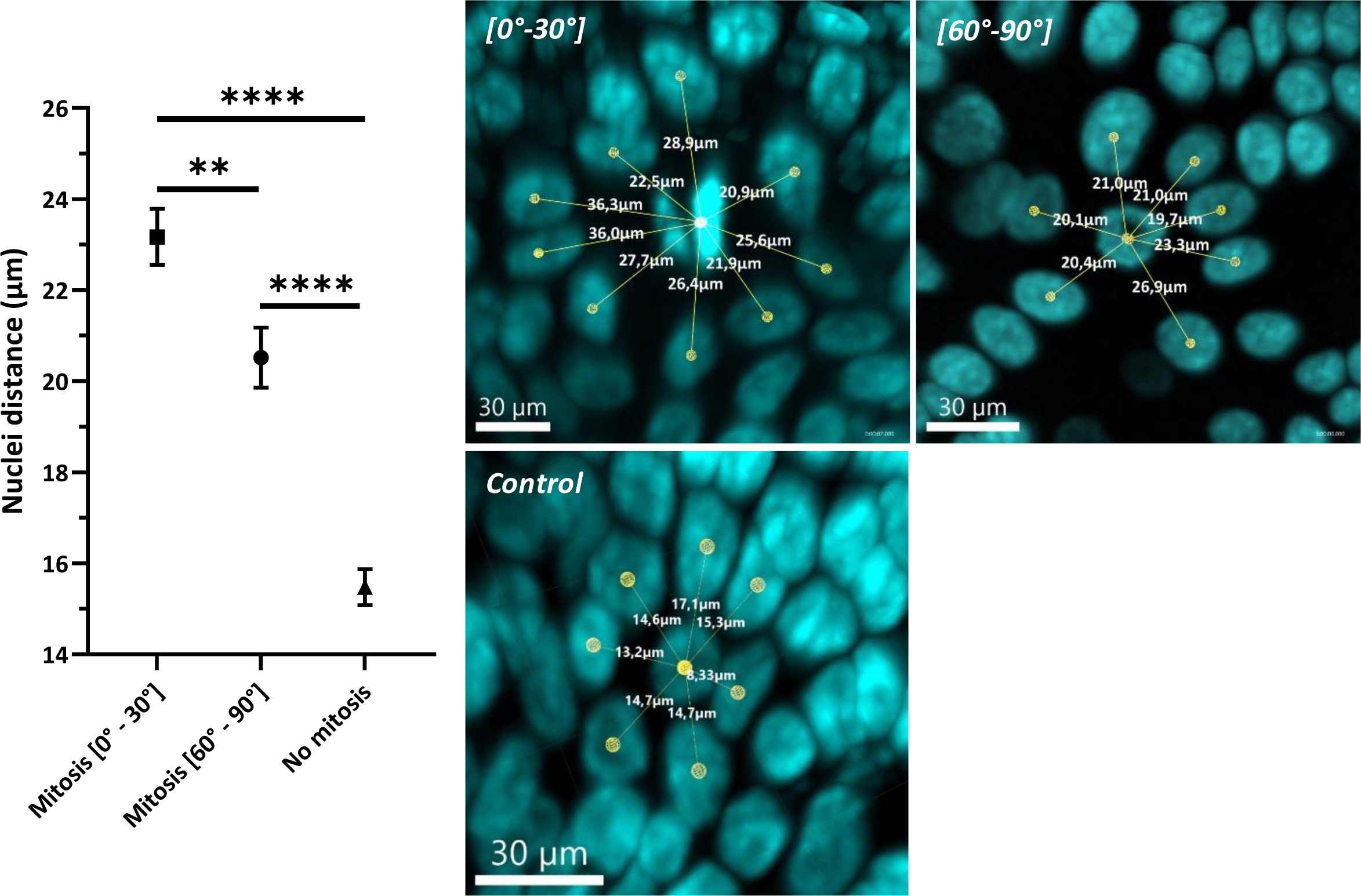
Measurements of the distances between the dividing nucleus and its surrounding nuclei. On the left, graph presenting the distances between the dividing nucleus in metaphase and the nuclei of neighbouring cells for each type of division and for controls (without mitosis), (Mann-Whitney test, the measurements have been done on 11 divisions of both categories picked at random among the 45 asymmetric and 28 symmetric divisions and observed on 8 organoids from 3 patients (a minimum of 2 organoids/patient and a maximum of 3 organoids/patients were analysed), p-value = 0,0022 between the two types of division and p-value < 0,0001 between each of these groups and the control one, mean ± SEM). On the right, representative images for each type of division and a condition ‘control’ without cell division, with the distances represented. Points for measuring distances are positioned at the barycentre of nuclei obtained through segmentation. A cell is considered as a neighbouring cell if its membrane is adjacent to the membrane of the dividing cell, visualized with the tubulin labelling.

Metaphase being the most easily identifiable stage of the mitosis in the images, we quantified the number of neighbouring cells directly surrounding the dividing cells and measured the distance separating the nuclei of the neighbouring cells from the cell at this step of the mitosis. The barycentre of the nuclei serves as reference point for the measurements. The measurements have been done on 11 divisions of both categories picked at random among the 45 asymmetric and 28 symmetric divisions and observed on 8 organoids from 3 patients (a minimum of 2 organoids/patient and a maximum of 3 organoids/patients were analysed).

Asymmetric and symmetric dividing cells have a mean of respectively 7,55 ± 0,37 (mean ± SEM) and 7,82 ± 0,55 neighbouring cells. No significant difference is thus observed (Mann Whitney t-test, p-value = 0,9394). In the absence of mitosis, we counted an average of 6,25 ± 0,28 neighbouring cells for a given cell, thus slightly fewer than in the presence of division (Mann Whitney t-test, p-value = 0,0285 between control group and [60°-90°] group, and p-value = 0.0150 between control group and [0°-30°] group).

Regarding the distances, our data showed that in the absence of cell division, nuclei are significantly closer together (Mann Whitney t-test, p-value < 0,0001 between control group and [60°-90°] group, and p-value < 0,0001 between control group and [0°-30°] group). Comparing the two types of division, in the case of asymmetric division, the nuclei of surrounding neighbour cells are significantly closer to the dividing cell (Mann-Whitney test, p-value = 0.0022) than in the case of symmetric division (figure 3). This result could suggest that the displacement of the matrix induced by asymmetric division could be less important than the one induced by symmetric division.

### Impact of asymmetric and symmetric divisions on the 3D matrix displacements

We then studied the displacements induced by both types of division on the matrix surrounding the epithelium. The signals obtained by confocal fluorescence microscopy, thanks to the cell labels (nuclei in blue, tubulin in green, figure 2a) and the beads inserted into the matrix (in red, figure 2a) to visualize the organoids and the ECM respectively, were processed in two distinct ways.

First, we established a framework that enables the tracking of the nuclei displacement over the three hours of culture (figure 4a): on the raw images, we first performed a 3D segmentation of the nuclei signal using Cellpose, and then used the Imaris software to track the nuclei over time. Indeed, Cellpose allows to train the segmentation model according to a machine-learning principle and, if necessary, to manually correct the segmentation. Segmentation using Cellpose therefore enables customizable segmentation with a high degree of precision, unlike segmentation using Imaris, where global segmentation parameters are selected and applied to all objects. This approach allowed us to perfectly perform the nuclei segmentation in 3D for our study, facilitating the follow-up of the nuclei displacement using Imaris.

**Figure 4.**
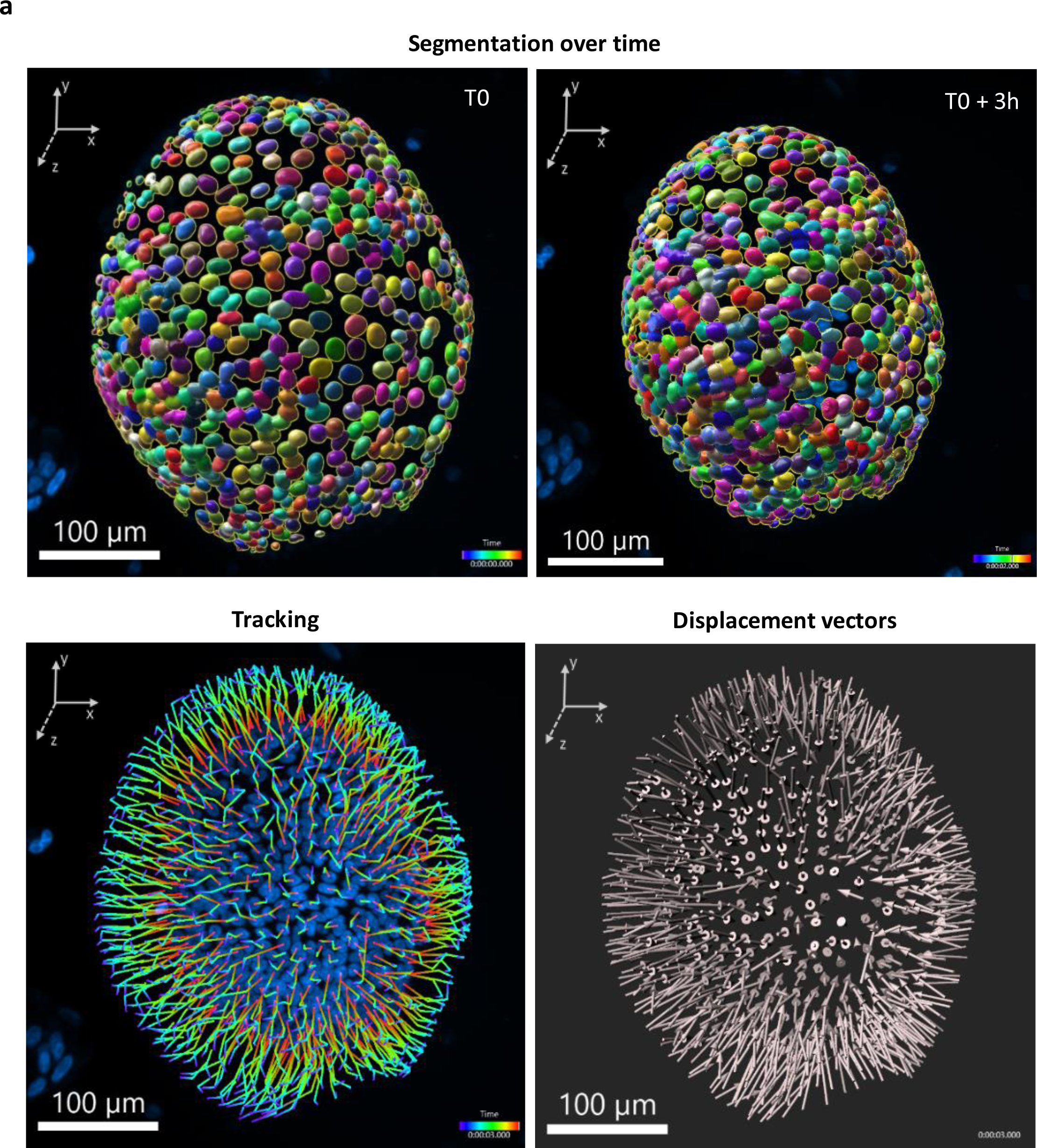
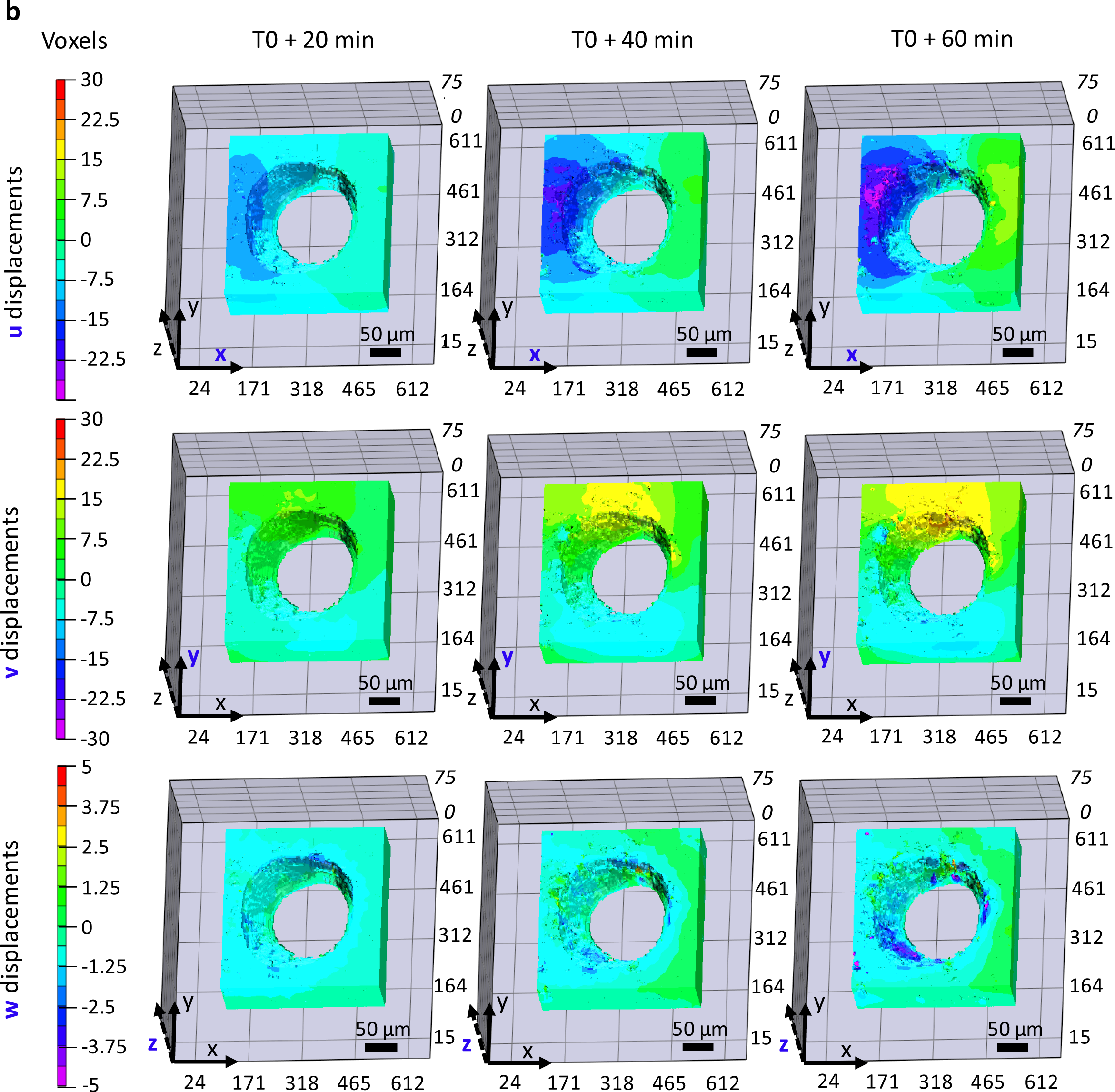
Displacements of organoid nuclei over time obtained after segmentation and tracking. (**a**) Nuclei are segmented and tracked over time, respectively with CellPose and Imaris. After the labelling and acquisition, nuclei are segmented with CellPose at each time point during 3h (Segmentation over time, visualisation of the segmentation masks in Imaris viewer) and tracked over time with Imaris tracker (Tracking). The global displacement (Displacement vectors) illustrates the uniform movement of nuclei. All scales are on the concerned images. (**b**) Displacement fields of the matrix obtained by digital volume correlation. Displacements of the matrix after 20 min (left), 40 min (middle) and 60 min (right). Voxel displacements in each axis are shown: u in the first row, v in the second row, w in the third row. The colour scale is in voxels. The voxel size is 0.64 x 0.64 x 2µm.

As shown in figure 4a and based on the measurements reported in table 1, the global analysis of nuclei displacements within the organoids enabled us to establish that, for a giving organoid, the nuclei display an overall uniform displacement as shown in a representative experiment (figure 4a).

**Table 1.**
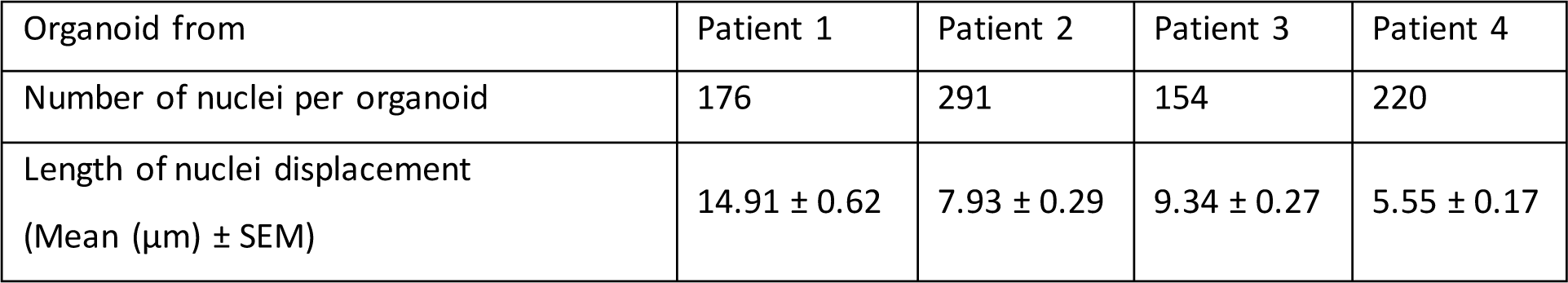
Nuclei displacements in organoids.

| Organoid from | Patient 1 | Patient 2 | Patient 3 | Patient 4 |
| --- | --- | --- | --- | --- |
| Number of nuclei per organoid | 176 | 291 | 154 | 220 |
| Length of nuclei displacement<br>(Mean ($\mu\text{m}$) ± SEM) | $14.91 \pm 0.62$ | $7.93 \pm 0.29$ | $9.34 \pm 0.27$ | $5.55 \pm 0.17$ |

Next, the signal allowing to follow the displacement of the matrix is reported by the tracking of the fluorescent beads dispersed within the Matrigel® between the time-points acquired every 20 minutes over the three hours of acquisition. It is used to perform image correlation, and more precisely DVC, using the VicVolume software. DVC is an experimental technique based on the use of two volumetric images (3D), one in a reference state, and the other in a deformed state. The principle of DVC is to slice the reference volumetric image using a grid to obtain sub-volumes (composed of voxels). Then, the correlation is performed to search individually each of the sub-volumes obtained from the reference image in the deformed image. The result is the displacement field of the material under study. According to our observations, mitosis is completed within one hour. In our analysis, we used as initial time (T0) the initiation of the mitosis (prophase) and performed the correlation between the four images taken every 20 minutes up to T0+60min once the mitosis is completed.

Dense volume correlation shows that the extracellular matrix undergoes global displacements, in each of the three spatial directions (figure 4b). However, unlike nuclei displacements, matrix displacement is not uniform. In line with our hypothesis, this suggests that specific localized biological events may affect the matrix differently, and in particular cell divisions and their orientation. Indeed, the colour map used, where green corresponds to zero displacement, clearly shows that the matrix is stressed as a whole, but not homogeneously. For example, from the first column onwards, we can see the appearance of a displacement along the x axis (“u displacements”), which propagates over time and gains in amplitude. In addition, we observed that nuclei displacements and matrix displacements do not occur at the same scale. Indeed, nuclei exhibit displacements of the order of ten micrometers (Table 1), while matrix displacements occur over greater distances. Indeed, as seen in figure 4b, the matrix undergoes a global displacement of up to several tens or even hundreds of micrometers around the organoid.

We postulated that local displacements of the matrix could correspond to cell division events and hypothesized that monitoring the 3D ECM displacements could be used as a read out for mitoses impact onto the matrix.

### Symmetric and asymmetric divisions impact the matrix differently

Penetrating the volume correlation along the z axis, we found that the presence of cell division generated a local displacement of the matrix, as exemplified in figure 5a. Indeed, on the right-hand side of the images, the green colour indicates a matrix displacement of 0 µm (so no displacement), while on the left-hand side we observed a wave of displacement whose intensity is strongest at the level of mitosis, identified in the first photo, and propagates throughout the (x,y) plane.

**Figure 5.**
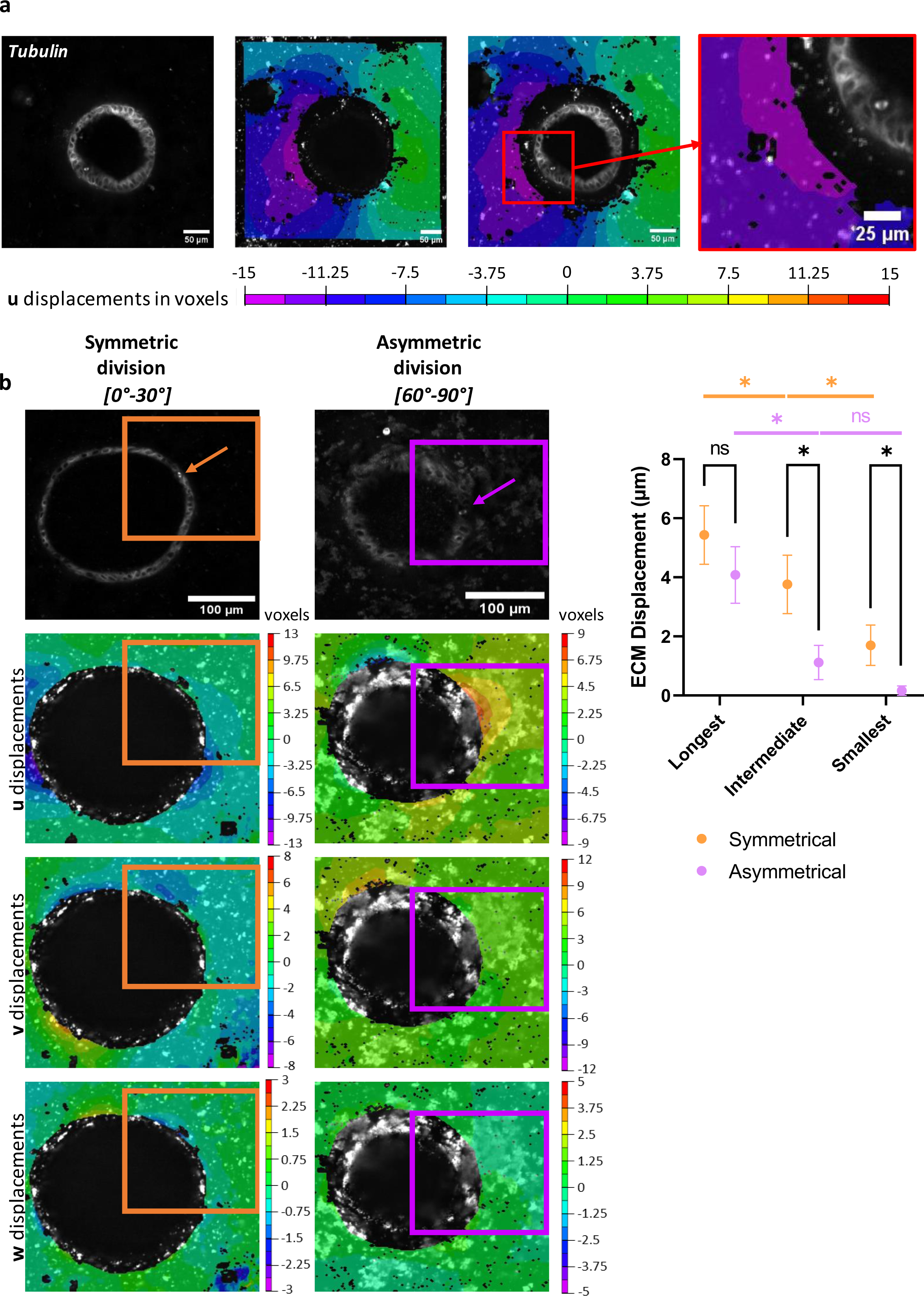
Impact of the mitotic spindle orientation on ECM. (**a**) Zoom on a z-axis section containing mitosis. From left to right: tubulin, ECM displacements along the u axis in the same section plane, merge of tubulin channel and matrix displacements, zoom on the area where cell division takes place. All scales are on the concerned images. (**b**) ECM displacements due to symmetric or asymmetric division. On the top panel, tubulin labelling showing the mitotic spindle. The arrows indicate the mitotic spindle. On the bottom panel, representation of matrix displacements on the mitosis z-slice, calculated in 3D, in the three axes. Orange and magenta squares represent the areas impacted by the mitosis. On the right, representation of the longest, intermediate and smallest displacements of the ECM for each type of cell division. Representation of the mean ± SEM (t test, 8 measurements in each group, p-value = 0.045 between both smallest displacements, p-value = 0.037 between intermediate displacements and p-value = 0.034 between longest displacements. t test between longest and intermediate displacement in asymmetric division: p-value = 0.0146; and in symmetric: p-value = 0.0152. t test between intermediate and smallest displacement in asymmetric divisions: p-value = 0.1109; and in symmetric one: p-value = 0.0474).

Having established that cell divisions participate in local ECM movements, we set out to determine the role of mitotic spindle orientation in these displacements. To solely investigate the impact of independent mitosis on matrix displacements and not of combined mitosis events, we only studied cases where mitosis was isolated, in both time and space. Here, 8 divisions of each type, observed in 5 organoids from 2 patients, were studied using these criteria. For each mitosis, matrix displacements in each of the three directions were measured at the cell-ECM frontier, and sorted relative to the two others under the labels of “longest displacement”, “intermediate displacement” and “smallest displacement” with values between 0 (i.e. no displacement) and 10 µm. For asymmetric and symmetric divisions, the longest displacement is significantly greater than the intermediate one (paired t test; asymmetric: p-value = 0.0146; symmetric: p-value = 0.0152). The intermediate displacement is significantly greater than the smallest one only for symmetric divisions (paired t test; asymmetric: p-value = 0.1109; symmetric: p-value = 0.0474) as shown in figure 5b.

Only 1 out of 8 asymmetric cell divisions causes the matrix to stain in the three directions, 2 cause a displacement in two directions while 5 cause a movement in the matrix in only one direction. In this case, the axis of matrix displacement corresponds to the axis of mitotic spindle orientation in the same plane. The longest displacement observed is an average of 4.08 ± 1.90 µm (mean ± SEM) for this type of mitosis. We can see that displacements in the other two directions are much smaller, averaging 1.11 ± 1.39 µm (mean ± SEM) for the intermediate displacement and 0.16 ± 0.28 µm (mean ± SEM) for the smallest displacement (figure 5b).

Symmetric cell divisions, on the other hand, always impact the matrix in several directions, either in two (3 out of 8 symmetric mitosis) or three directions (5 out of 8). In fact, the mean of the longest displacements is 5.44µm ± 2.31 µm (mean ± SEM), slightly greater than for asymmetric divisions, but without any significant difference. On the other hand, the mean of the intermediate displacement is 3.76 ± 2.21 µm (mean ± SEM). Similarly, the smallest displacement is 1.7 ± 1.48 µm (mean ± SEM) (figure 4b).

From a qualitative standpoint, we can conclude that asymmetric divisions involve a rather uniaxial displacement, whereas symmetric mitoses cause a multiaxial one. In addition, from a quantitative standpoint, the displacements induced in the case of symmetric division are systematically greater. Considering the longest displacement, there is no significant difference between the one observed in symmetric divisions and the one in asymmetric division (respectively of 5.44µm ± 2.31 and 4.08 ± 1.90 µm, unpaired t test, p-value = 0.3421). Nevertheless, considering the intermediate displacement, we can note that it is 3 times greater in the case of symmetric divisions than the one observed in asymmetric divisions (3.76 ± 2.21 and 1.11 ± 1.39 µm, unpaired t test, p-value = 0.0373). At last, we observed a factor of 10 between the smallest displacements of symmetric and asymmetric cell divisions (1.7 ± 1.48 and 0.16 ± 0.28 µm, unpaired t test, p-value = 0.0453).

## Discussion

The study of mechanical interactions between the epithelium and its matrix, particularly the impact of cell division in real time and in 3D, remains complex to this day due to technical constraints^10^. By choosing the human colon organoid, we ensured the suitability of the biological model on some major criteria allowing this type of study: physiological relevance (different types of cell division), rapid growth and the accessibility of the ECM. In order to study the ECM displacements, we developed a DVC approach, never used before for studying ECM displacements in a three-dimensional environment *in vitro*. In this article, we have thus provided a new method for measuring and analysing the mechanical interactions between the colonic epithelium and its ECM. In fine, this approach allowed us to visualize and analyse over time in 3D the displacements of both the organoid and its surrounding matrix. We observed a strong overall contraction of the epithelium overtime. The displacements of the matrix obtained by DVC over a period of one hour correspond to a contraction following the “u” direction and an expansion following the “v” direction. Analysis of the positions of neighbouring cells during mitoses indicates greater proximity during the asymmetrical divisions than during the symmetrical divisions. Moreover, the analyses carried out during the cell division phases show greater movements of the matrix during symmetrical mitoses than during asymmetrical division.

Previous studies have highlighted the ability of stem cells and early progenitors to divide respectively symmetrically and asymmetrically which is crucial in adapting the needs of the colon during homeostasis, growth, or regeneration ^14,16^. Each type of division is involved in distinct processes of intestinal homeostasis: symmetric mitosis are responsible for epithelial renewal, while asymmetric ones are involved in the differentiation process ^16^. However, while a number of factors that can orient direct cell division are well known ^22–25^, how spindle orientation affects the matrix environment remains largely unexplored. Our results show that asymmetric divisions appear to have a local and uniaxial impact on the matrix, in contrast to the symmetric divisions that affect the matrix more widely in different axis. From a biological perspective, this finding could be of interest to be further study in a pathological context. As an example, it could provide clues to the formation of bifid crypts, one of the first visible sign of tumour initiation, which rely on area of multiple hyperproliferative cells within the crypt. As changes in ECM composition, rigidity and/or organization may activate pro-tumour signalling pathways, it is possible that matrix remodelling due to symmetric division participate to this process ^26^. In addition, it is well known that intestinal stem cells can increase the proportion of symmetric mitosis in response to environmental cues. In order to assess the mechanical interactions between the epithelium and its matrix in a cancer context, it may therefore be interesting to work with tumour organoids.

## Methods

### Human colon organoids

The organoids used in this study are part of the registered COLIC biobank (DC-2015-2443) (for details, refer to ^27^). The culture was established from colonic samples obtained from surgical resections of patients treated for colorectal cancer at the Toulouse University Hospital that gave informed consent. Tissues were harvested in healthy zones at the margin of the resection and at least 10 cm away from the tumour. Organoids are stored in liquid nitrogen to ensure optimum preservation and are thawed when needed. After thawing, organoids undergo a passage: 3D structures are dissociated using trypsin (*TrypLE*®, Invitrogen) supplemented with 10 μM Y27632 during 10 min at 37°C. Mechanical agitation takes place every 3 minutes to promote dissociation. Digestion is stopped by adding washing medium (DMEM/F12 supplemented with 5% SFV and 10μM of Y27632) at 4°C. The solution containing dissociated organoids is then centrifuged 5 min at 400g, at 4°C. The cell pellet is recovered and re-suspended in Matrigel®. Matrigel® domes are deposited in 96-well culture plates (10µL/well) with glass bottom. Next, 100µL of pro-stemness culture medium ^28^ is added to each well, and renewed every two days. Cell growth occurs in an incubator at 37°C and 5% CO_2_, during 10 days.

### Fluorescent Beads Adding

Fluorescent silica beads are added into the Matrigel® to obtain a speckle pattern (10% v/v Matrigel® beads) just before the Matrigel® dome formation stage. The beads (Sicastar Red-F, Micromod) are 1µm-diameter silica beads covered with a red fluorophore (excitation 569 nm / emission 585 nm).

### Fluorescent probes

3h prior microscope image acquisition, organoid culture medium is removed and replaced by fresh medium containing the fluorescent probes to stain the DNA (Hoechst, 1:3000 v/v) and the tubulin (Biotracker 488 Microtubules, Merck, dilution 1:500 used with Verapamil 1:1000 to contain the probe inside the cells).

### Imaging Set-up

At day 10 of culture, the organoids are imaged in a thermostatically-controlled chamber (37°C), buffered with CO_2_ (5%) with the Opera Phenix™ microscope (Perkin Elmer). The objective used is a x20 water objective. The z-step used is 2 µm, covering a z-depth of around 100 µm (i.e. around 50 z-slice). Images are acquired every 20 minutes during 3 hours.

### Image pre-processing

The images of the beads are processed using the “Basic” plugin on ImageJ, which enables illumination defects to be corrected in terms of depth. This step is required to obtain a uniform signal that can be used for DVC. The nuclear signal is denoised with the “Subtract Background” option on ImageJ.

### Mitotic orientation measurements

The angular orientations correspond to the angle between mitotic spindle and ECM at the basal side of the cell. The measurements of mitotic spindle orientations are made in the acquisition plane (x,y), and normalized relative to the basal side of the organoid in ImageJ. The angles thus obtained are classified into 3 categories, based on the literature ^17^: [0-30°], [30-60°] and [60-90°].

### Nuclei segmentation

Nuclei were segmented using the Cellpose 2.0 machine-learning-based approach. The system is trained with nuclei images from several organoids. The training images are 2D slice, taken in all the possible plans (x,y), (x,z) and (y,z) ^29^. After the training, z-stack are segmented plan by plan and then the 2D segmentations were assembled in 3D using the same software to obtain 3D labels of nuclei.

### Nuclei tracking

The segmentation mask obtained with CellPose is used in the commercial Imaris software to track the nuclei over time. The barycentre of each nucleus is used to perform the tracking thanks to the “Surface” Tool of Imaris.

### Matrix displacement analysis

Digital volume correlation (DVC) was performed using VicVolume software (*Correlated Solutions®)* on fluorescent beads images. A subset size between of 29 voxels and a step of 3 are optimal. As illustrated in figure 6, the image z-stack made up of voxels is digitally sliced using a grid of user-selected size, adapted to the size and density of the speckle pattern. Each window thus created (orange square) on the reference image, i.e. the first time point “t”, is then searched for in the other z-stacks, corresponding to the deformed states, i.e. the subsequent time points “t +Δt”. By obtaining the translational and rotational displacements for each window, a displacement field map is obtained for the entire image. Matrix displacements were thus analysed in 3D, along each of the three axes corresponding to the three dimensions. The reference frame used is (x,y,z), where (x,y) corresponds to the microscope acquisition plane. Displacements in u correspond to matrix displacements along the x axis. Similarly, v displacements correspond to displacements along the y axis, and w displacements to displacements along the z axis. The size of a voxel is (0.64, 0.64, 2) µm.

**Figure 6.**
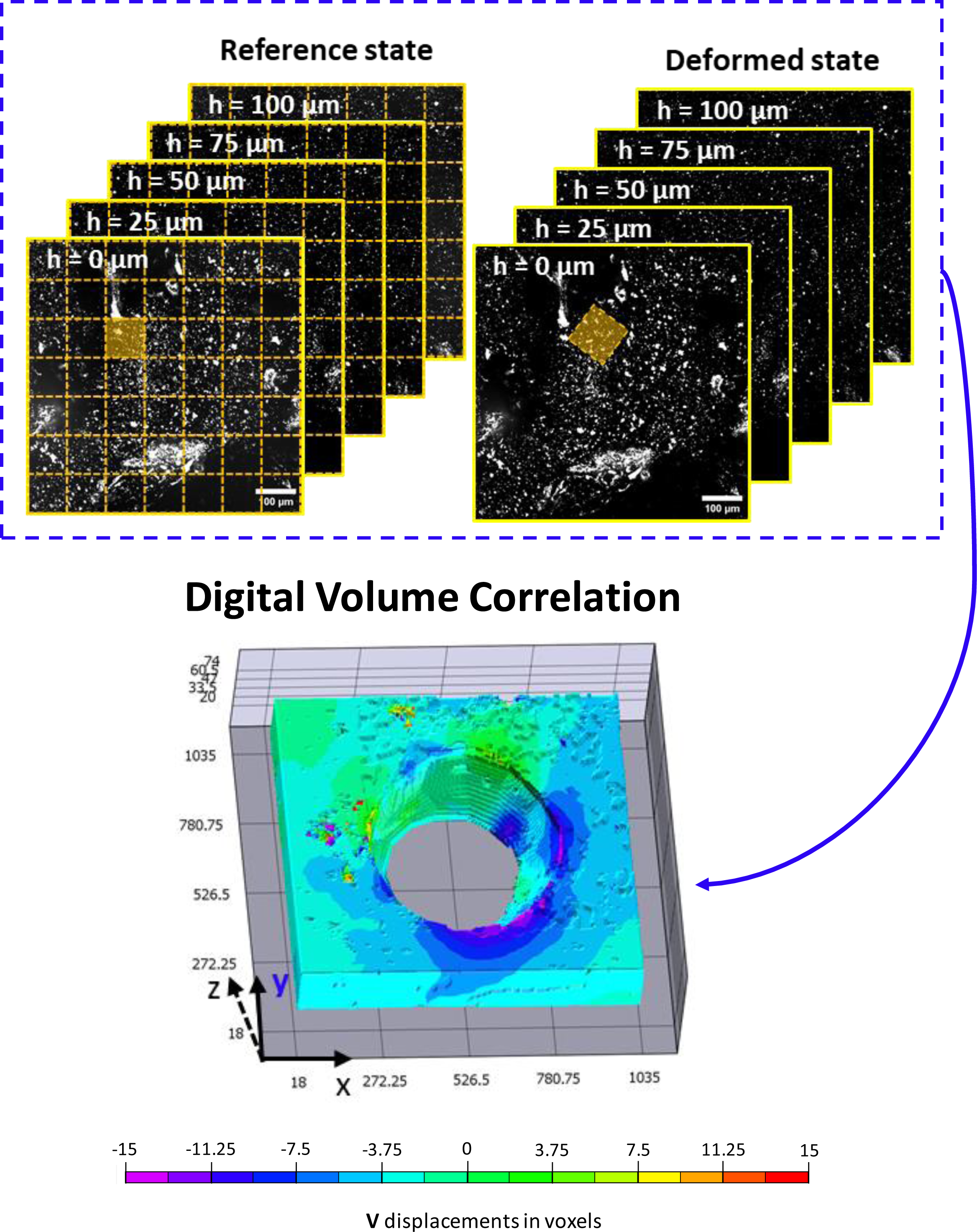
Digital Volume Correlation (DVC) principle. DVC is performed on the beads stack images thanks to VicVolume to obtain ECM displacements fields between a reference state and a deformed state of the matrix.

### Statistical analysis

Data are expressed as mean ± SEM. Analyses were performed using the GraphPad Prism 9 software. Statistical significance was determined by paired or unpaired t-test or Mann Whitney test, the use of one or the other is mentioned in the text. Significance was accepted for p<0.05 for each test performed.

## Data availability

The data used and/or analyzed during this study are available from the corresponding author on reasonable request.

## Acknowledgements

This collaborative work was funded by two Cancer Plan projects: Mocassin - Biosystem 2017, granted to AF and Melchior - MIC 2020, granted to FB. JL is supported by both. DM is supported by Plan cancer Melchior-MIC 2020. LM is supported by the INSERM and the Région Occitanie. We also wish to thank the patients who agree in giving their tissue for research purposes and allowed this study, and Camille Douillet for critical discussions and suggestions on this work. We thank David Sagnat for technical assistance at the organoids facility of Inserm UMR 1220, Toulouse, France.

## Author contributions

Audrey Ferrand, Florian Bugarin, Stephane Segonds and Gaelle Recher organized the project. Léa Magne, Déborah Michel and Julien Laussu collected the data. Léa Magne and Thomas Pottier performed data analysis and visualization. Léa Magne, Audrey Ferrand and Florian Bugarin drafted the manuscript. Audrey Ferrand and Florian Bugarin led the project and oversaw manuscript preparation. All authors have read and approve the submitted manuscript.

## References

1. Gupta, V. K. et al. The nature of cell division forces in epithelial monolayers. Journal of Cell Biology 220, e202011106 (2021).

2. Barker, N. et al. Identification of stem cells in small intestine and colon by marker gene Lgr5. Nature 449, 1003–1007 (2007).

3. Medema, J. P. & Vermeulen, L. Microenvironmental regulation of stem cells in intestinal homeostasis and cancer. Nature 474, 318–326 (2011).

4. Onfroy-Roy, L., Hamel, D., Foncy, J., Malaquin, L. & Ferrand, A. Extracellular Matrix Mechanical Properties and Regulation of the Intestinal Stem Cells: When Mechanics Control Fate. Cells 9, 2629 (2020).

5. Lu, P., Takai, K., Weaver, V. M. & Werb, Z. Extracellular Matrix Degradation and Remodeling in Development and Disease. Cold Spring Harb Perspect Biol 3, a005058 (2011).

6. Lussier, C., Basora, N., Bouatrouss, Y. & Beaulieu, J.-F. Integrins as mediators of epithelial cell-matrix interactions in the human small intestinal mucosa. Microscopy Research and Technique 51, 169–178 (2000).

7. Katsumi, A., Orr, A. W., Tzima, E. & Schwartz, M. A. Integrins in Mechanotransduction *. Journal of Biological Chemistry 279, 12001–12004 (2004).

8. He, H. et al. Piezo channels in the intestinal tract. Front Physiol 15, 1356317 (2024).

9. Orr, A. W., Helmke, B. P., Blackman, B. R. & Schwartz, M. A. Mechanisms of Mechanotransduction. Developmental Cell 10, 11–20 (2006).

10. Pérez-González, C., Ceada, G., Matejčić, M. & Trepat, X. Digesting the mechanobiology of the intestinal epithelium. Current Opinion in Genetics & Development 72, 82–90 (2022).

11. Pérez-González, C. et al. Mechanical compartmentalization of the intestinal organoid enables crypt folding and collective cell migration. Nat Cell Biol 23, 745–757 (2021).

12. He, S. et al. Stiffness Restricts the Stemness of the Intestinal Stem Cells and Skews Their Differentiation Toward Goblet Cells. Gastroenterology 164, 1137–1151.e15 (2023).

13. Guevara-Garcia, A., Soleilhac, M., Minc, N. & Delacour, D. Regulation and functions of cell division in the intestinal tissue. Seminars in Cell & Developmental Biology 150–151, 3–14 (2023).

14. Shahriyari, L. & Komarova, N. L. Symmetric vs. Asymmetric Stem Cell Divisions: An Adaptation against Cancer? PLoS One 8, e76195 (2013).

15. Snippert, H. J. et al. Intestinal crypt homeostasis results from neutral competition between symmetrically dividing Lgr5 stem cells. Cell 143, 134–144 (2010).

16. Joly, A. & Rousset, R. Tissue Adaptation to Environmental Cues by Symmetric and Asymmetric Division Modes of Intestinal Stem Cells. Int J Mol Sci 21, 6362 (2020).

17. Quyn, A. J. et al. Spindle Orientation Bias in Gut Epithelial Stem Cell Compartments Is Lost in Precancerous Tissue. Cell Stem Cell 6, 175–181 (2010).

18. Knoblich, J. A. Asymmetric cell division during animal development. Nat Rev Mol Cell Biol 2, 11– 20 (2001).

19. Nakajima, Y. Mitotic spindle orientation in epithelial homeostasis and plasticity. The Journal of Biochemistry 164, 277–284 (2018).

20. Lechler, T. & Mapelli, M. Spindle positioning and its impact on vertebrate tissue architecture and cell fate. Nat Rev Mol Cell Biol 22, 691–708 (2021).

21. Spence, J. R. et al. Directed differentiation of human pluripotent stem cells into intestinal tissue in vitro. Nature 470, 105–109 (2011).

22. Fink, J. et al. External forces control mitotic spindle positioning. Nat Cell Biol 13, 771–778 (2011).

23. Petridou, N. I. & Skourides, P. A. FAK transduces extracellular forces that orient the mitotic spindle and control tissue morphogenesis. Nat Commun 5, 5240 (2014).

24. Xie, J. et al. Contribution of cytoplasm viscoelastic properties to mitotic spindle positioning. Proc Natl Acad Sci U S A 119, e2115593119 (2022).

25. Saleh, J. et al. Length limitation of astral microtubules orients cell divisions in murine intestinal crypts. Developmental Cell 58, 1519–1533.e6 (2023).

26. Brauchle, E. et al. Biomechanical and biomolecular characterization of extracellular matrix structures in human colon carcinomas. Matrix Biol 68–69, 180–193 (2018).

27. Sébert, M. et al. Thrombin modifies growth, proliferation and apoptosis of human colon organoids: a protease-activated receptor 1-and protease-activated receptor 4-dependent mechanism. Br J Pharmacol 175, 3656–3668 (2018).

28. Sato, T. et al. Long-term expansion of epithelial organoids from human colon, adenoma, adenocarcinoma, and Barrett’s epithelium. Gastroenterology 141, 1762–1772 (2011).

29. Pachitariu, M. & Stringer, C. Cellpose 2.0: how to train your own model. Nat Methods 19, 1634– 1641 (2022).

